# Investigation of region-specific effects of pepsin-digested decellularized meniscus on human adipose-derived stromal cells within alginate hydrogels

**DOI:** 10.1101/2025.09.17.676824

**Authors:** Sheradan Doherty, Xindi Zhao, Anna Kornmuller, Pascal Morissette Martin, Lauren E. Flynn

## Abstract

Meniscus tears are one of the most common musculoskeletal injuries, but treatment options remain limited. The meniscus can be divided into distinct inner and outer regions based on differences in the tissue structure and composition. Recognizing that the extracellular matrix (ECM) provides cell-instructive cues that can direct cell differentiation, the current study investigated the effects of incorporating region-specific ECM derived from decellularized meniscus on the viability, growth, and lineage-specific differentiation of human adipose-derived stromal cells (ASCs) encapsulated in alginate beads. The first phase of research focused on validating a novel decellularization protocol for porcine meniscus, which demonstrated the effective removal of cellular content while preserving key ECM constituents including glycosaminoglycans (GAGs) in both the inner and outer meniscus regions. Subsequently, the decellularized inner meniscus (DIM) and decellularized outer meniscus (DOM) were digested with pepsin and combined with a propriety alginate formulation to enable rapid cell encapsulation under mild conditions. Viability staining confirmed that encapsulated human ASCs remained highly viable over 28 days in culture in proliferation or chondrogenic differentiation media. Under both media conditions, the ASC density was significantly higher in the alginate beads incorporating DIM or DOM as compared to alginate alone controls at 28 days. In addition, gene expression analysis and immunofluorescence staining supported that the incorporation of the pepsin-digested ECM within the alginate enhanced fibrochondrogenic marker expression in the samples cultured in chondrogenic differentiation medium. Qualitatively, more intense staining for collagen types I and II were observed within the beads incorporating DOM as compared to DIM, supporting that the formulations had varying effects. Overall, these studies provide new insight supporting that region-specific meniscus ECM can be harnessed to direct cell phenotype and function towards the goal of developing tissue-specific bioinks for tissue-engineering applications.

## 1. Introduction

The menisci are a pair of crescent-shaped fibrocartilaginous tissues that play important roles in shock absorption, load transmission, joint stability, and joint lubrication in the knee [1,2]. Both the medial and lateral meniscus exhibit a unique zonal organization and biochemical composition that gives rise to distinct inner and outer regions. Specifically, the inner region of the meniscus resembles cartilage tissue, including a more abundant population of chondrocyte-like cells, as well as an extracellular matrix (ECM) that is enriched in collagen type II and glycosaminoglycans (GAGs) [3]. In contrast, the outer region is more ligament-like, containing fibroblast-like cells and a higher abundance of collagen type I [3,4].

The meniscus is the most frequently injured structure in the knee joint and with a limited capacity for self-repair, meniscal injuries are one of the most common orthopaedic conditions requiring surgical intervention [3,5,6]. While meniscal repair can be effective for young patients, success rates depend heavily on the vascularization of the meniscal zones. Degenerative tears in the avascular inner region are often treated by meniscectomy, involving partial or complete removal of the meniscus [3,7]. These procedures have been shown to alter the biomechanics of the joint, ultimately accelerating the onset of osteoarthritis (OA) [1,8]. As such, there is a critical need for novel approaches targeting meniscus repair.

Notably, tissue-engineering strategies have shown promise in stimulating meniscal regeneration [9]. Previous studies have explored the use of naturally-derived [10] and synthetic [11] implants, as well as the injection of pro-regenerative cell populations such as bone marrow- or adipose-derived mesenchymal stromal cells (MSCs) into the injured regions [12]. Recognizing the cell-instructive capacity of the ECM, there is evidence to support that the incorporation of tissue-specific ECM sourced from decellularized tissues within hydrogel delivery vehicles could help to promote MSC-mediated tissue regeneration by providing biological cues that direct cell migration, proliferation, and differentiation [13]

To date, the majority of meniscal decellularization protocols include treatment with sodium dodecyl sulphate (SDS) [14–17], an anionic detergent that is associated with ECM denaturation and cytotoxicity, which may compromise ECM bioactivity [18]. Recognizing these limitations, our first step was to develop a new decellularization protocol for porcine meniscus using the non-ionic detergent Triton X-100, which has been reported to better preserve ECM composition and structure in other tissue types compared to SDS treatment [19].

Following validation of successful decellularization of both the inner and outer meniscus regions, we sought to integrate the decellularized meniscus into a hydrogel platform for controlled cell encapsulation that would be compatible with future bioprinting applications. We selected alginate as the hydrogel carrier as it offers advantages in terms of its mild and rapid gelation conditions, cell-supportive nature, and low cost [20]. The decellularized meniscus was enzymatically-digested using pepsin to generate a peptide solution that could be readily incorporated within the alginate. Pepsin digestion is a common approach for ECM hydrogel fabrication, and previous studies have demonstrated that pepsin-digested ECM can have cell-instructive effects in mediating cell proliferation and lineage-specific differentiation [13,21].

We hypothesized that the incorporation of the pepsin-digested decellularized meniscus within the alginate would enhance the fibrochondrogenic differentiation of encapsulated MSCs *in vitro*.

Building from previous work that demonstrated that decellularized meniscus has region-specific effects on the differentiation of rabbit synovium-derived MSCs [22], we sought to compare hydrogels incorporating pepsin-digested ECM sourced from decellularized inner meniscus (DIM) versus decellularized outer meniscus (DOM) to alginate alone controls. Human adipose-derived stromal cells (ASCs) were selected as an accessible, clinically-relevant MSC source with the capacity to differentiate towards the chondrogenic lineage, which has shown promise in stimulating meniscal regeneration in pre-clinical models [23,24]. In-depth *in vitro* studies were performed to characterize ASC viability and retention, as well as differentiation towards a fibrochondrocyte-like phenotype under both proliferation and chondrogenic differentiation conditions to assess potential chondro-inductive and chondro-conductive effects of the region-specific ECM.

## 2. Methods

### 2.1 Materials

Unless otherwise indicated, all chemical reagents were purchased from Sigma-Aldrich Canada Ltd. (Oakville, Ontario).

### 2.2 Meniscal decellularization

Meniscus samples were harvested from freshly-isolated knee joints of pigs (5-8 months in age) sourced from the Mount Brydges Abattoir (Mount Brydges, ON, Canada). Menisci were cut along the midline to separate the inner and outer regions, which were processed separately for all studies. The tissues were cut into 2.0 mm segments using a biopsy punch to increase the surface area. Unless otherwise noted, all incubations were performed at 37 °C under agitation at 120 rpm, and all solutions were supplemented with 1% antibiotic/antimycotic solution, as well as phenylmethylsulfonyl fluoride (PMSF) for the first 3 days of processing.

One gram of tissue was subjected to two freeze-thaw cycles (−80 °C / 37 °C) in 50 mL of hypotonic buffer (pH 8.0) composed of 10 mM tris (hydroxymethyl)aminomethane (Tris) and 5 mM ethylenediaminetetraacetic acid (EDTA) in deionized water (dH_2_O), with solution changes between each cycle, followed by an additional 24 h incubation with a solution change after the first 8 h. Next, the samples were subjected to a high-salt detergent extraction (pH 8.0) in 50 mL of 50 mM Tris buffer supplemented with 1% (v/v) Triton X-100 and 1.5 M KCl for 24 hours, with a solution change after 12 h. The samples were then rinsed three times for 30 minutes in 50 mL of Sorenson’s phosphate buffer (SPB) rinsing solution comprising 0.55 M sodium phosphate dibasic heptahydrate (Na_2_HPO_4_·7H_2_O) and 0.17 M potassium phosphate (KH_2_PO_4_) (pH 8.0). Next, the samples were enzymatically digested for 2 hours in 50 mL of SPB digest solution composed of 0.55 M Na_2_HPO_4_·7H_2_O, 0.17 M KH_2_PO_4_, and 0.049 M magnesium sulphate heptahydrate (MgSO_4_·7H_2_O) (pH 7.3) supplemented with 300 U/mL DNase Type II and 20 U/mL RNase Type III. The samples were then subjected to a final detergent extraction in 1% (v/v) Triton X-100 in 50 mM Tris buffer (pH 8.0) overnight for 18 hours. Finally, the detergent solution was replaced in the morning and the samples were incubated for an additional 8 hours. At the end of the process, the samples were rinsed three times for 30 min in dH_2_O, frozen at −80°C in dH_2_O, and lyophilized using a Labconco Freezone 4.5 lyophilizer (Labconco, Kansas City, MO, United States) for 48 h.

### 2.3 Histological analyses of native and decellularized tissues

Native and decellularized meniscus samples (n=3, N=3) were rehydrated in phosphate-buffered saline (PBS) and fixed overnight at 4°C in 4% paraformaldehyde (PFA). The samples were embedded in paraffin and sectioned (7 μm sections) for staining with toluidine blue to visualize GAG content, picrosirius red to visualize collagen, and 4’,6-diamidino-2-phenylindole (DAPI) in fluoroshield mounting medium (Abcam) to visualize cell nuclei, following standard protocols. Images were obtained using an EVOS XL Core microscope, a Nikon Optiphot polarizing microscope, and an EVOS FL fluorescence microscope, respectively.

### 2.4 Scanning electron microscopy

Scanning electron microscopy (SEM) was performed on native and decellularized meniscus samples to visualize the ECM ultrastructure (n=3, N=2). Briefly, lyophilized tissues were thinly sectioned and coated with osmium. Samples were imaged with a LEO 1530 scanning electron microscope at an accelerating voltage of 1 kV and a working distance of 3.6-3.8 mm.

### 2.5 Biochemical analyses of native and decellularized tissues

For biochemical analyses, lyophilized decellularized meniscus samples and native tissue controls were cryo-milled using a Retsch Mixer Mill MM 400 milling system. The cryomilled samples were digested in 500 µL of Tris-EDTA (TE) buffer supplemented with 600 U Proteinase K (Qiagen) for 4 h at 56 °C, with vortexing every 15 min (n=3 separately digested samples/batch, N=3 decellularization batches for all assays).

A Quant-iT™ PicoGreen® assay (Molecular Probes) was used to quantify the dsDNA content in the native and decellularized tissues. Briefly, DNA was extracted from the digested samples using the DNeasy Blood & Tissue Kit (Qiagen), following the manufacturer’s instructions. The native tissue samples were diluted 1:15 in TE buffer, while the decellularized samples were processed undiluted. The samples were combined with the Quant-iT™ reagent within black 96-well plates and the fluorescence was read using a CLARIOstar^®^ microplate reader. The dsDNA concentration in each sample was determined through comparison to a lambda DNA standard curve and was normalized based on dry weight.

The dimethylmethylene blue (DMMB) assay was used to quantify the sulphated glycosaminoglycan (sGAG) content in the digested samples. In brief, the samples were diluted 1:20 in PBS supplemented with 1 w/v% bovine serum albumin (BSA) and 10 µL of the diluted sample was combined with 200 µL of DMMB reagent (0.016 mg/mL in 0.2% formic acid (pH 5.3) in 96-well plates. The absorbance was read using a CLARIOstar^®^ microplate reader at 525 nm. The GAG concentration of each sample was determined using a chondroitin sulphate standard curve and was normalized based on dry weight.

### 2.6 Immunohistochemical analyses of native and decellularized tissues

Lyophilized native and decellularized meniscus samples (n=3, N=3) were rehydrated through an ethanol series (100%, 90%, 70%, 50% (v/v) diluted in PBS) for 30 min each and rinsed twice in PBS for 15 min at room temperature. Samples were then embedded in Tissue-Tek OCT compound and snap frozen in liquid nitrogen before cryosectioning (7 μm sections). Following fixation in acetone for 10 minutes at −20°C and blocking in 10% goat serum in tris-buffered saline with 0.1% tween (TBST), sections were stained overnight at 4°C with primary antibodies against collagen type I (dilution 1:100 in TBST with 2% BSA, ab34710, Abcam), collagen type II (dilution 1:200, ab34712, Abcam), collagen type IV (dilution 1:100, ab6586, Abcam), collagen type VI (dilution 1:300, ab6588, Abcam), fibronectin (dilution 1:150, ab23750, Abcam), and laminin (dilution 1:200, ab11575, Abcam). Detection was carried out using an anti-rabbit secondary conjugated to Alexa Fluor 594 (dilution 1:200, ab150080, Abcam) or an anti-mouse secondary conjugated to Alexa Fluor 650 (dilution 1:200, ab96882, Abcam). No primary antibody controls were included in all trials. Images were acquired with an EVOS FL fluorescence microscope.

### 2.7 Adipose-derived stromal cell isolation and culture

Subcutaneous adipose tissue samples were collected with informed consent from female patients undergoing elective lipo-reduction surgery in London, Ontario, with approval from the Human Research Ethics Board at Western University (HSREB# 105426). The tissue samples were transported to the lab in sterile PBS supplemented with 20 mg/mL BSA on ice and immediately processed to isolate human ASCs following published methods [25]. ASCs were grown on tissue culture polystyrene flasks in proliferation medium (DMEM/Ham’s F12 supplemented with 10% fetal bovine serum (FBS) (Gibco®, Invitrogen, Burlington, ON, Canada) and 100 U/mL of penicillin with 0.1 mg/mL streptomycin (pen-strep) (Gibco®, Invitrogen, Burlington, ON, Canada)) and passaged at 80% confluence. All experiments were performed with passage 4 (P4) ASCs.

### 2.8 Pepsin digestion of cryomilled meniscus

Cryomilled DIM and DOM, processed separately, were added at a concentration of 20 mg/mL to sterile 10% (w/w) porcine pepsin (3200-4500 mU/mg protein) in 0.1 M hydrochloric acid and digested for 18 h at 37 °C under agitation at 120 rpm. The solutions were neutralized with sterile 1 M sodium hydroxide and undigested fragments were removed by filtering through a sterile 40 μm cell strainer. The resultant solutions were stored at 4 °C for a maximum of 48 h before use.

### 2.9 ASC encapsulation in alginate beads

A propriety alginate formulation (AG-10 Matrix, Aspect Biosystems Ltd., Vancouver, British Columbia) was combined in a 1:1 ratio with pepsin-digested DIM (ALG+DIM) or DOM (ALG+DOM) to obtain a final ECM concentration of 10 mg/mL and gently mixed with ASCs at a density of 3.0 x 10^6^ cells/mL. The beads were drop-casted through a 16 G needle into a bath of calcium-based crosslinker (CAT-2, Aspect Biosystems Ltd., Vancouver, BC, Canada). Pure alginate beads (ALG) were also synthesized as controls. The beads were incubated in the crosslinking solution for 10 minutes at 37 °C. The resultant beads were transferred to 12-well culture inserts (Greiner Bio-one) and rinsed once with proliferation medium.

Bead diameter was measured using a previously established method [26], involving supplementation of the CAT-2 crosslinker with 0.05% (w/v) Coomassie Brilliant Blue for visualization purposes. Immediately following crosslinking, the beads (n=3, N=3) were imaged at 4X magnification using an EVOS XL Core microscope and analyzed using ImageJ analysis software.

### 2.10 Culture of encapsulated ASCs

Following ASC encapsulation, the beads were cultured in proliferation medium at 37 °C, 5% CO_2_. After 72 h, the proliferation medium was replaced in the induced set of samples with chondrogenic differentiation medium [27] composed of proliferation medium supplemented with 10 ng/mL recombinant human transforming growth factor-β1 (TGF-β1) (BioLegend), 50 μg/mL ascorbate-2-phosphate, 6.25 μg/mL bovine insulin, and 100 nM dexamethasone. Non-induced samples maintained in proliferation medium were included in all assays to assess potential chondro-inductive effects of the ECM peptides.

### 2.11 Analysis of ASC viability and density

The viability and density of ASCs within the alginate beads were assessed through confocal microscopy in both induced and non-induced samples collected at 24 h, 7 days, 14 days and 28 days post-encapsulation, using the LIVE/DEAD^®^ Assay (Invitrogen CAT#L3224) (n=3 beads from each composition/timepoint, N=3 cell donors). The beads were incubated in 4 μM ethidium homodimer-1 (EthD-1) and 2 μM calcein-AM in PBS for 30 min and imaged using the 5X objective at 3 depths separated by 40 μm on a Zeiss LSM800 Confocal Microscope with Airyscan. ImageJ analysis software was used to count the viable cells to compare the ASC density within the beads at 24 h and 28 days post-encapsulation.

### 2.12 RT-qPCR analysis of chondrogenic gene expression

At 7 and 28 days post-encapsulation (i.e. days 4 and 25 post-induction of chondrogenic differentiation for the induced samples), RT-qPCR analysis of chondrogenic gene expression was performed. The ASCs were released from the beads through incubation in 25 mM sodium citrate in dH_2_O for 45 min at 37 °C under agitation at 120 rpm. Total RNA was extracted in PureZol (Bio-Rad Laboratories) and purified using the Aurum Total RNA Fatty and Fibrous Tissue kit (Bio-Rad Laboratories), according to the manufacturer’s instructions. cDNA was synthesized using the iScript™ cDNA Synthesis Kit (Bio-Rad Laboratories) from 300 ng of input RNA in a 20 μL volume. Gene expression was analyzed by real-time qPCR using the BioRad CFX-384 system using 2xSSoAdvancedTM Universal SYBR® Green Supermix (Bio-Rad Laboratories). The panel of markers is summarized in Table 1, with glucuronidase beta (*GUSB*) and ribosomal protein L 13a (*RPL13A)* included as stable housekeeping genes. The cycling protocol was: 95 °C for 2 min, followed by denaturation at 95 °C for 10 s and annealing and elongation at 60 °C for 30 s, repeated for 40 cycles. Transcript levels were analyzed using the ΔΔC_T_ method, with normalization of each sample to the geometric mean of the housekeeping genes and using the induced ALG samples at 7 days post-encapsulation as the calibrator (n=3 samples processed separately at each timepoint/trial, N=2 trials with different ASC donors).

**Table 1.**
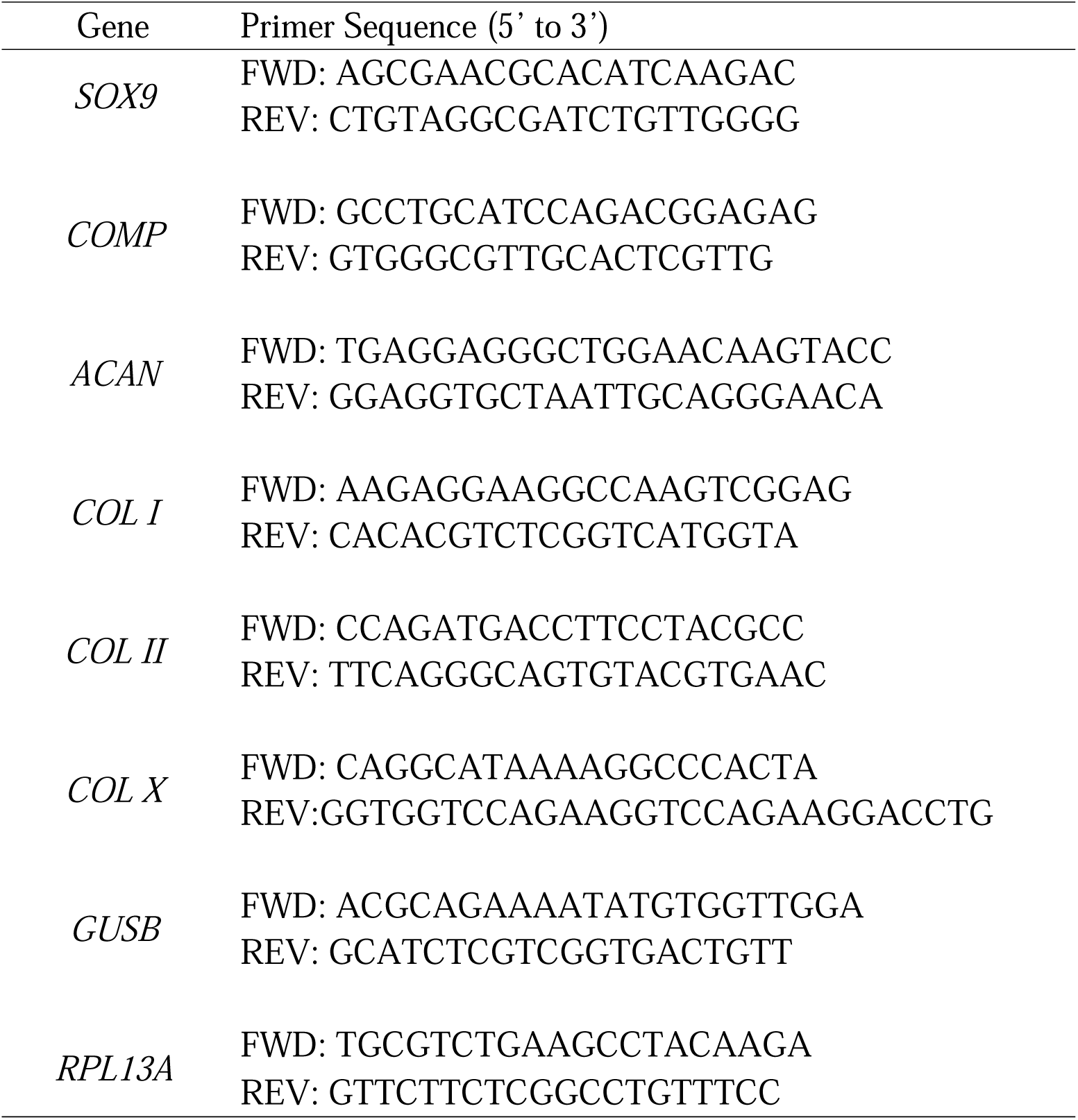
RT-qPCR Primer Sequences.

### 2.13 Immunofluorescence analysis of chondrogenic ECM marker expression

Immunofluorescence staining was performed to qualitatively assess the presence of key chondrogenic ECM components in the ALG+DIM, ALG+DOM and ALG alone beads at 28 days post-encapsulation. Briefly, beads were collected at each timepoint and fixed in 4% paraformaldehyde with 10 mM calcium chloride and 0.1 M sodium cacodylate for 4 h at 4°C. The samples were then washed in 0.1 M sodium cacodylate with 50 mM barium chloride for 18 h at 4°C prior to paraffin embedding and sectioning (7 μm sections). Sections were deparaffinized by rinsing in xylene (two times for 5 minutes each) and rehydrated in an ethanol series (100%, 100%, 95%, 70%), followed by dH_2_O, for 2 minutes each.

Heat-mediated antigen retrieval was performed using DAKO Target Retrieval Solution (DAKO Canada Inc., Burlington, ON, Canada). Next, the samples were fixed with acetone for 10 min at −20°C, and blocked with 10% goat serum in TBST, prior to overnight incubation at 4°C with the primary antibodies listed above against collagen type I (dilution 1:100 in TBST with 10% goat serum), collagen type II (dilution 1:400), and fibronectin (dilution 1:150). Detection was carried out using an anti-rabbit secondary conjugated to Alexa Fluor 594 (dilution 1:200, ab150080, Abcam) or an anti-mouse secondary conjugated to Alexa Fluor 650 (dilution 1:200, ab96882, Abcam) (n=3 cross-sections containing multiple beads/trial, N=2 trials with different ASC donors). No primary antibody controls were included in all trials. Images were acquired using an EVOS FL fluorescence microscope.

### 2.14 Statistical Analyses

All numerical data are expressed as mean ± standard deviation (SD). All statistical analyses were performed by two-way ANOVA with a Tukey’s post-hoc comparison of the means using GraphPad Prism 7 (GraphPad Software, San Diego, CA). Differences of p < 0.05 were considered statistically significant.

## 3. Results

### 3.1 New meniscus decellularization protocol effectively extracts cellular content while preserving key ECM components

The first goal of this project was to establish a decellularization protocol not involving SDS treatment that would effectively remove cellular content from both the inner and outer regions of porcine menisci, while retaining the complex ECM composition. The inner and outer regions were processed separately to enable the investigation of potential region-specific cell-instructive effects of the processed ECM on the survival and fibrochondrogenic differentiation of human ASCs. Qualitative analysis of cell nuclei in the processed samples through DAPI staining showed no detectable nuclei remaining in the decellularized tissues from both regions at the end of processing (Figure 1(a)). Effective decellularization was confirmed through quantitative analysis via the PicoGreen® assay, which showed significant reductions (∼99%) in dsDNA content compared to native tissue samples from both regions (Figure 1(b)).

**Figure 1.**
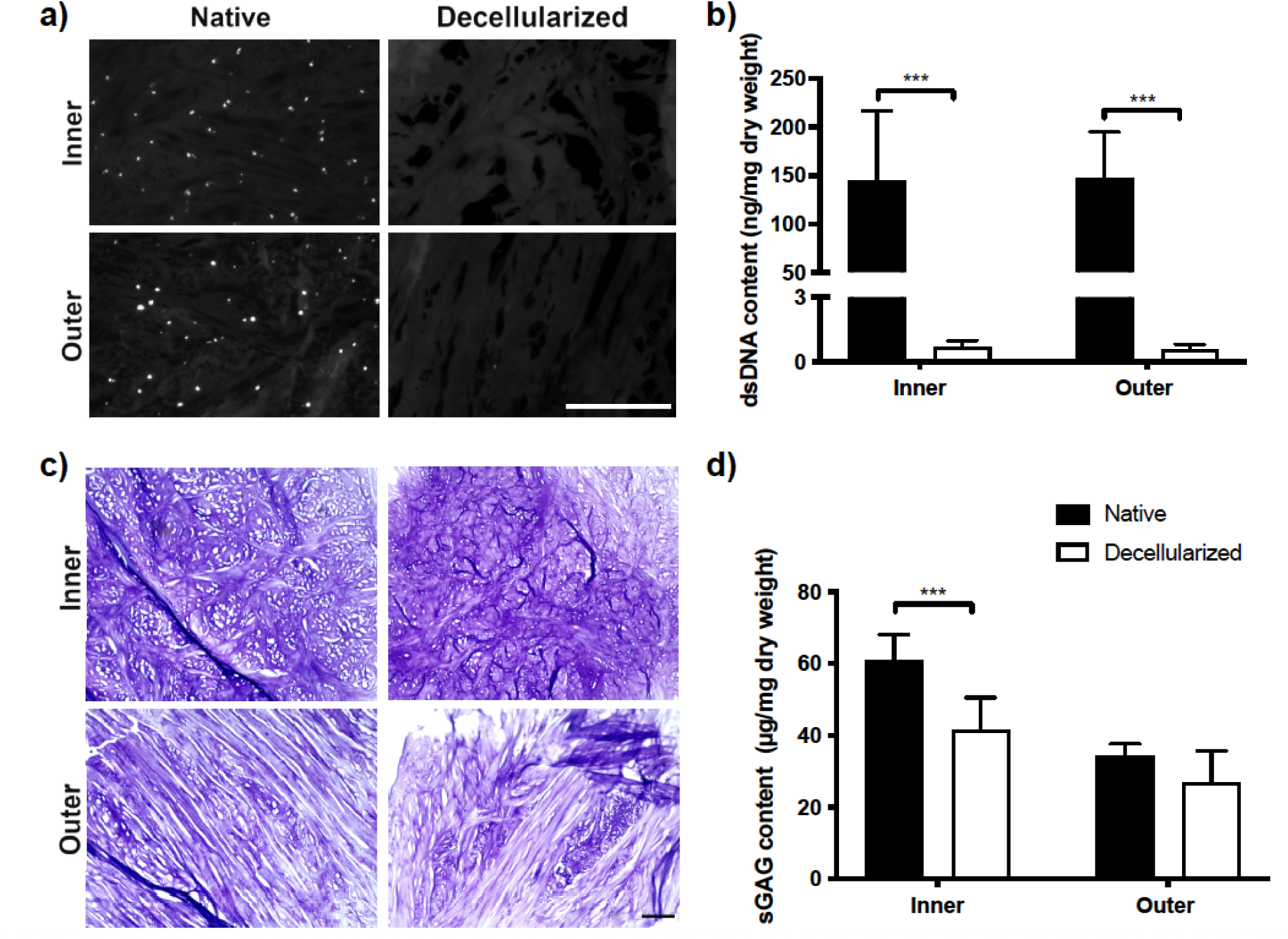
New decellularization protocol involving treatment with Triton-X100 was effective at decellularizing porcine menisci. **(a)** Representative DAPI nuclear staining of decellularized inner and outer meniscus samples and native tissue controls showing no detectable cell nuclei (white) following decellularization. Scale bar=200 μm. (**b)** Quantitative analysis of dsDNA content using the PicoGreen assay confirms effective cell removal from both regions. **(c)** Representative toluidine blue staining showing retention of GAGs (purple) following decellularization. Scale bar=200 μm. **(d)** Quantitative analysis of sGAG content via the DMMB assay, showing a significant decrease in sGAG content in the inner region following decellularization. (n=3, N=3), ***p<0.0001.

Toluidine blue staining qualitatively confirmed that GAGs were retained in the samples from both regions following decellularization (Figure 1(c)). Quantitative analysis of sGAG content via the DMMB assay revealed a significant loss of sGAG from the inner region, with ∼68% retained following decellularization. In contrast, there was no significant difference in sGAG content between the native and decellularized samples from the outer region, with ∼79% of sGAG retained (Figure 1(d)). Picrosirius red staining revealed dense networks of collagen, including a combination of thicker (red-orange) and thinner (yellow-green) fibres, with qualitatively similar patterns in the native and decellularized tissue samples (Supplemental Figure 1).

Based on SEM analysis, all samples had a complex fibrous ECM ultrastructure, with preservation of the collagen fibrillar banding patterns, and no obvious differences in the ECM between the native and decellularized samples (Figure 2).

**Figure 2.**
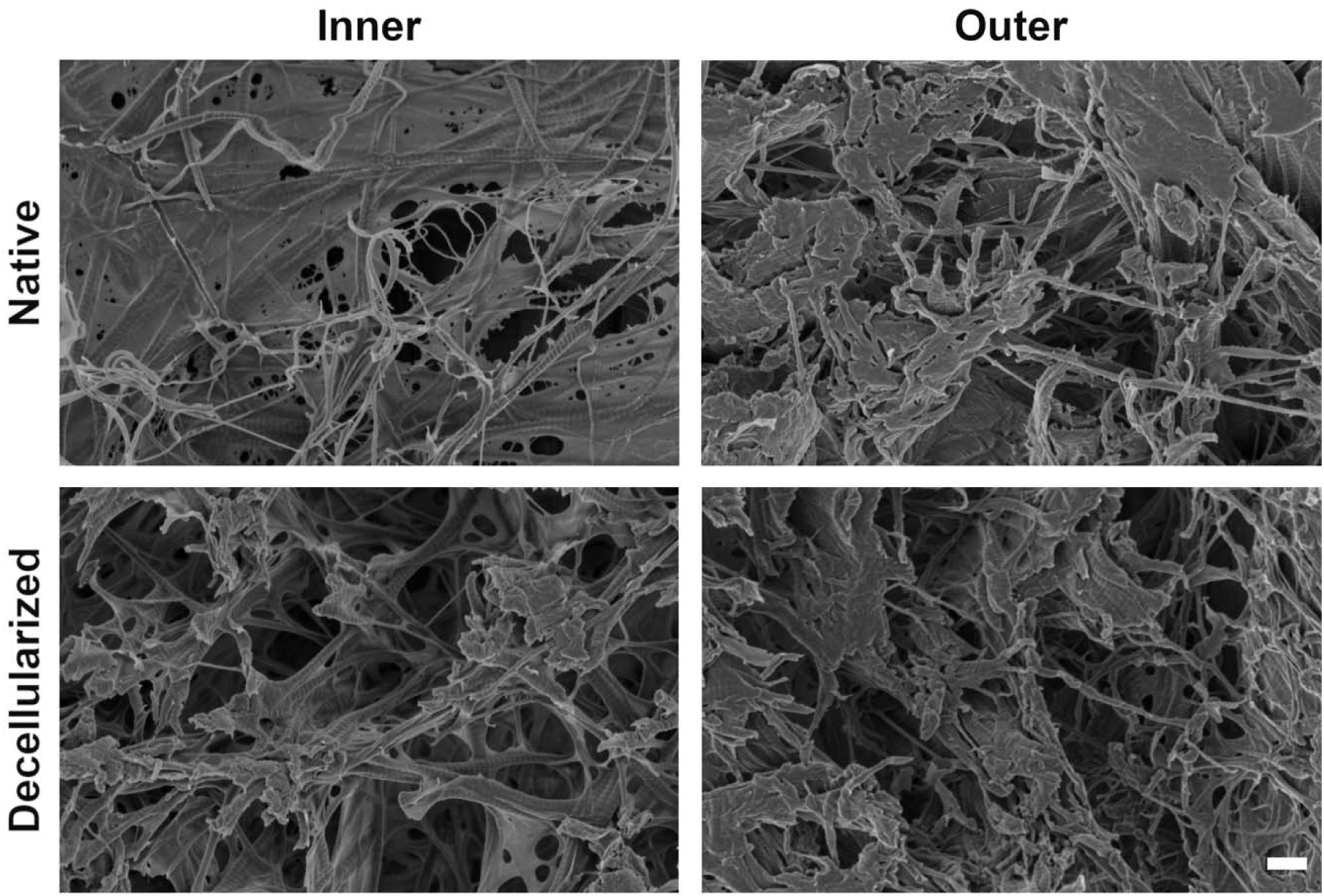
Representative SEM images of native and decellularized meniscus showing that all samples had a complex fibrous ECM ultrastructure. Scale bar=400 nm. Immunofluorescence staining was subsequently performed to characterize the presence and distribution of key meniscal ECM components in the inner and outer regions following decellularization relative to native tissue controls (Figure 3). Staining for collagen types I, II, IV, VI, fibronectin, and laminin showed qualitatively similar patterns between the native and decellularized meniscus samples in both regions. Consistent with the expected regional variations, collagen I and fibronectin staining appeared qualitatively more intense in the outer meniscus samples.

**Figure 3.**
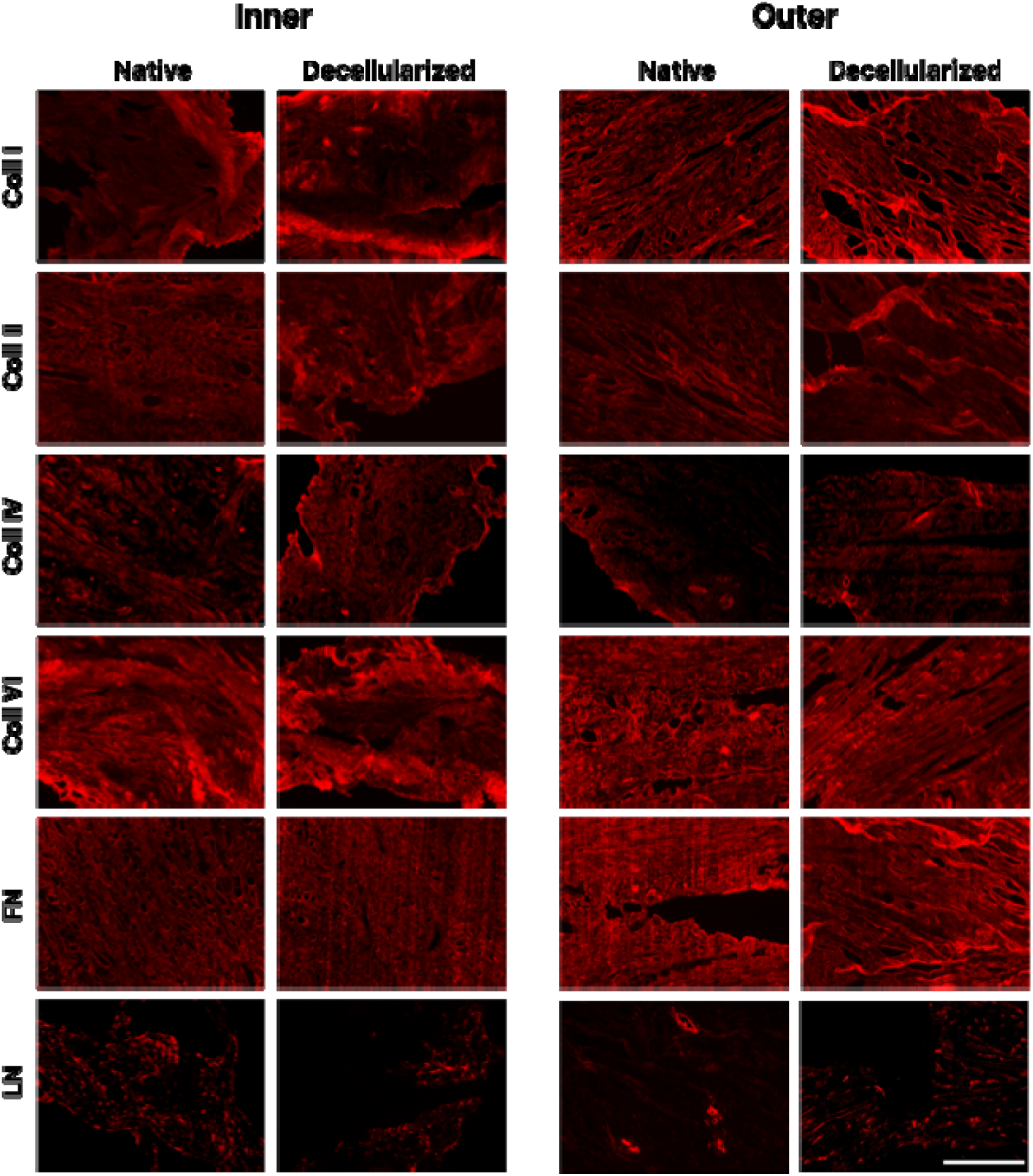
Immunohistochemical analyses confirmed the presence of key ECM components following decellularization. Representative staining showed similar patterns between native and decellularized meniscus for all ECM markers investigated. Scale bar=400 μm. Abbreviations: Coll I=collagen type I, Coll II=collagen type II, Coll IV=collagen type IV, Coll VI=collagen type VI, FN=fibronectin, LN=laminin.

### 3.2 Composite alginate-ECM beads support human ASC viability and retention

DIM and DOM samples were lyophilized, cryomilled, and enzymatically digested using pepsin. The resultant solubilized ECM was combined with a proprietary alginate formulation to generate region-specific hydrogel beads (ALG+DIM and ALG+DOM) containing a final ECM concentration of 10 mg/mL, based on the initial dry mass of the decellularized tissues within the digests. In addition, alginate (ALG) alone samples were included as a control. Each hydrogel formulation was combined with human ASCs at a density of 3.0 × 10^6^ cells/mL and dropcast into the propriety crosslinker to form beads. The average bead size for all three formulations was similar, with diameters measuring 3.8 ± 0.5 mm, 3.7 ± 0.4 mm, and 3.3 ± 0.4 mm for the ALG, ALG+DIM, and ALG+DOM beads respectively.

LIVE/DEAD^®^ staining with confocal imaging was performed to assess ASC viability following encapsulation and culture within the ALG, ALG+DIM, and ALG+DOM beads at 24 hours, 7 days, 14 days, and 28 days post-encapsulation and culture in proliferation medium (Figure 4(a) & Supplemental Figure 2) or chondrogenic differentiation medium (Figure 4(c) & Supplemental Figure 3). In all conditions, the encapsulated ASCs remained highly viable throughout the culture period. Quantification of the viable ASC density at 24 hours and 28 days post-encapsulation showed there was a significant increase in ASC density within the ALG+DIM group from 24 hours to 28 days in the samples cultured in proliferation medium (Figure 4(a)). In both media formulations, the ASC density within the ALG+DIM and ALG+DOM beads was significantly higher than in the ALG alone beads at 28 days (Figures 4(a), (b)).

**Figure 4.**
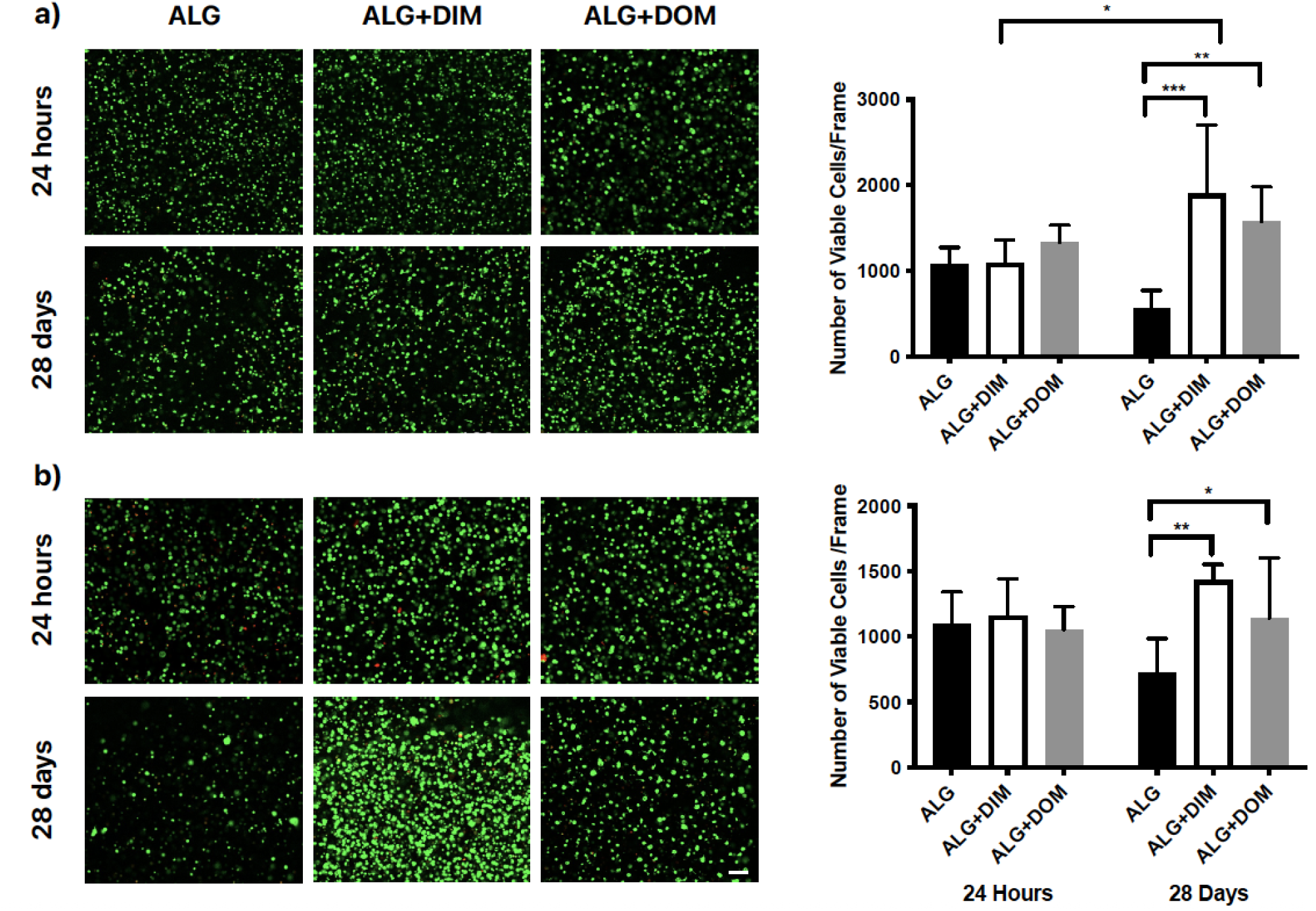
LIVE/DEAD staining confirmed that the human ASCs encapsulated in the alginate beads remained highly viable over 28 days in culture in proliferation and chondrogenic differentiation media. Representative images showing calcein^+^ live (green) and EthD-1^+^ dead (red) ASCs in the ALG, ALG+DIM, and ALG+DOM beads cultured in **(a)** proliferation medium or **(b)** chondrogenic differentiation medium. Scale bar = 100 μm. The samples were encapsulated and cultured for 3 days in proliferation medium, prior to being transferred into chondrogenic differentiation medium. Graphs on the right show live cell density, reported as cells/frame at 5X magnification. The density was significantly higher at 28 days in the ALG+DIM and ALG+DOM beads as compared to the ALG beads in both media conditions. For the samples cultured in proliferation medium, there was a significant increase in the ASC density in ALG+DIM beads from 24 hours to 28 days. (n=3, N=3), *p<0.05, **p<0.001, ***p<0.0001.

### 3.3 ECM incorporation enhanced chondrogenic gene expression in the encapsulated ASCs cultured in differentiation medium

RT-qPCR analysis was performed as an initial screen to analyze the effects of region-specific ECM incorporation on the expression of fibrochondrogenic genes in the encapsulated human ASCs at 7 and 28 days post-encapsulation. In assessing the samples cultured in proliferation medium, many of the genes were not consistently expressed, suggesting that there was a low-level of differentiation. The only notable pattern was that expression of *SOX9* and *Coll I* were more highly expressed at 7 days compared to 28 days in all groups (Supplemental Figure 4). For the samples cultured in chondrogenic differentiation medium (Figure 5), there was increased expression of *SOX9, COMP* and *ACAN* at 7 days post-encapsulation in the ALG+DOM beads relative to the other two formulations. Consistent with a progression in differentiation, expression of the ECM markers *ACAN, Coll I*, *Coll II,* and *Coll X* was generally higher after 28 days of chondrogenic culture compared to 7 days in all groups. Further, in comparing the expression levels in the 28-day chondrogenic samples, *ACAN* and *Coll II* were more highly expressed in the ALG+DOM beads relative to the ALG+DIM and ALG alone beads. In contrast, the hypertrophic marker *Coll X* was most highly expressed in the ALG+DIM beads.

**Figure 5.**
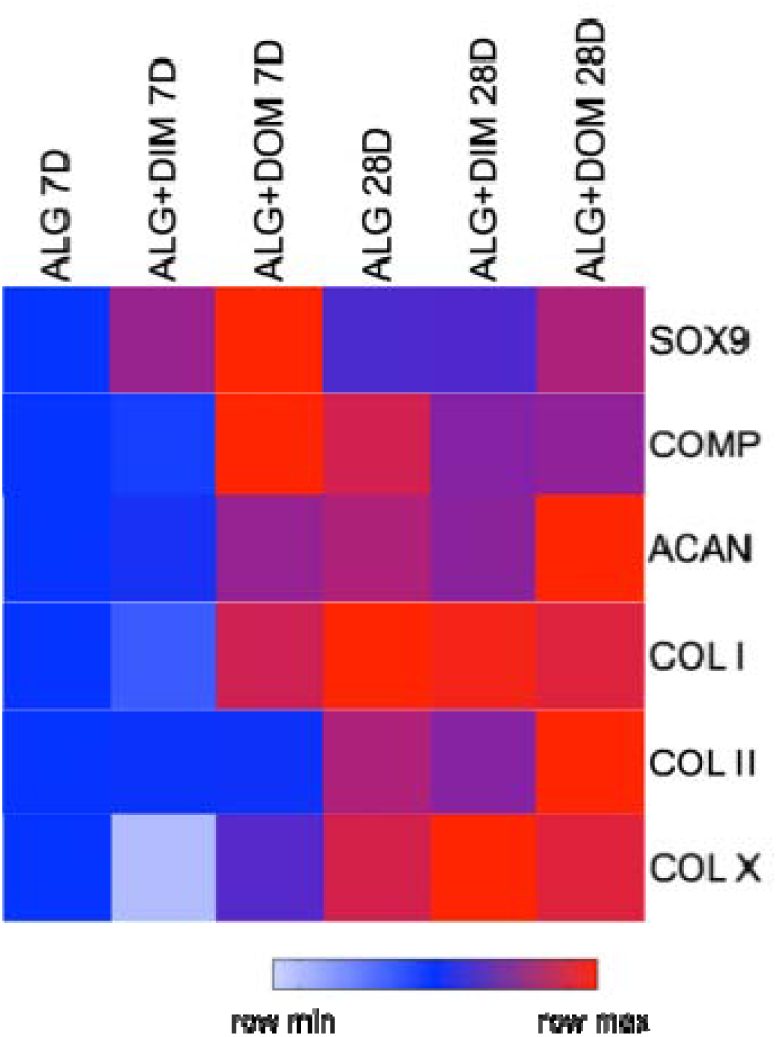
Fibrochondrogenic gene expression was enhanced in the human ASCs encapsulated within the alginate-based beads following 28 days of culture under chondrogenic conditions. The samples were encapsulated and cultured for 3 days in proliferation medium, prior to being transferred into chondrogenic differentiation medium for an additional 25 days of culture. Data was analyzed using the ΔΔCt method with normalization to the geometric mean of the stable housekeeping genes *RPL13A* and *GUSB*, and using the day 7 alginate beads as the calibrator. The heat map represents the average relative gene expression for the two cell donors, with similar patterns observed between the donors. (n=3 bead samples processed separately at each timepoint/trial, N=2 trials with different ASC donors).

### 3.4 Incorporation of pepsin-digested DIM and DOM enhanced fibrocartilaginous ECM accumulation in the samples cultured in chondrogenic differentiation medium

Immunofluorescence staining was performed to visualize collagen I, collagen II and fibronectin expression within the alginate-based beads at 28 days post-encapsulation. There was limited expression of the chondrogenic ECM markers detected in the beads cultured in proliferation medium, with the exception of fibronectin, which appeared to be qualitatively more highly expressed in the ALG+DIM beads (Supplemental Figure 5). While only low levels of the markers were detected in the ALG beads cultured in chondrogenic medium, markedly higher expression levels were observed in the induced ALG+DIM and ALG+DOM beads, with qualitatively higher levels of expression of collagen I and II in the ALG+DOM beads relative to ALG+DIM beads (Figure 6).

**Figure 6.**
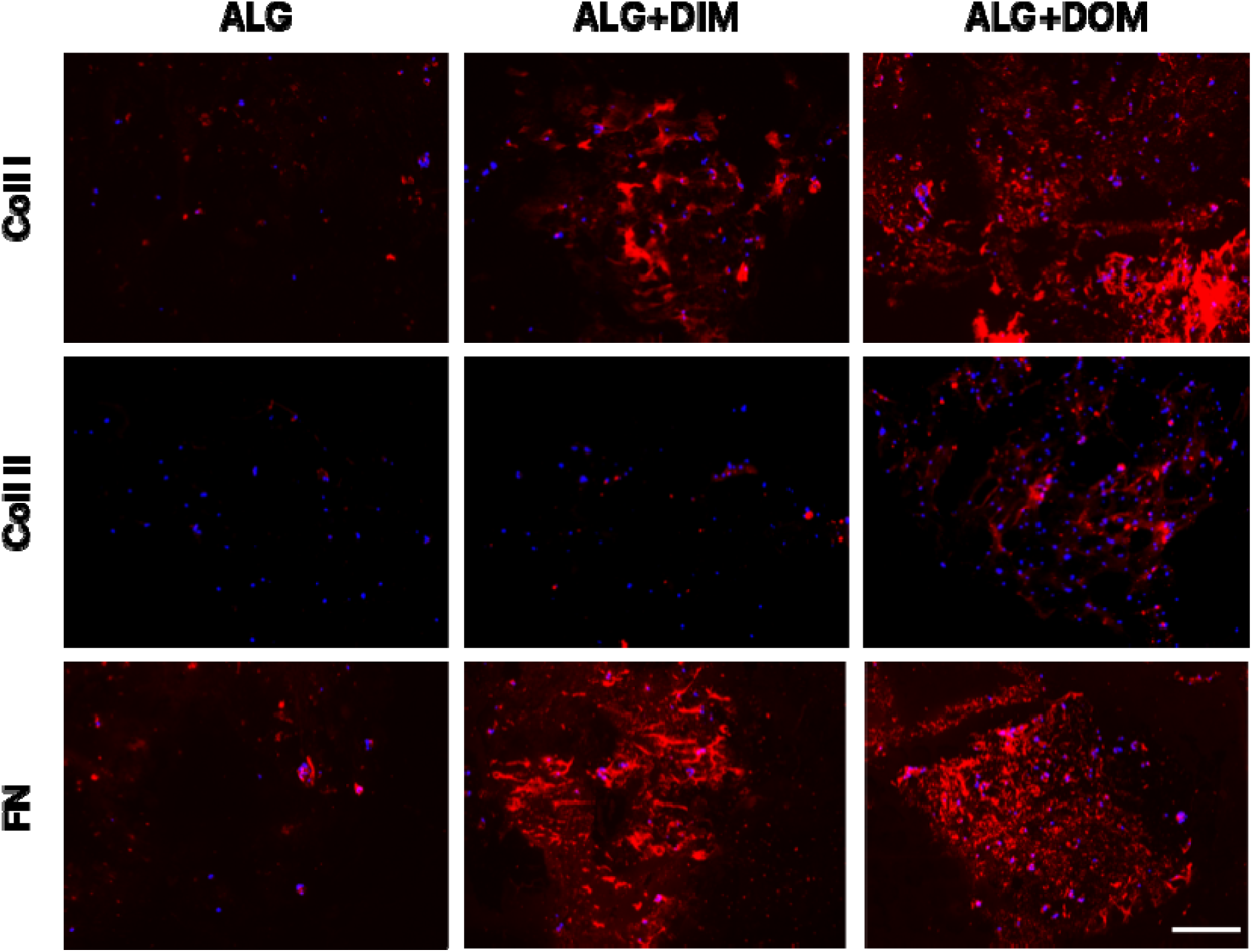
Immunofluorescence staining showing enhanced ECM marker expression within the composite alginate + ECM beads at 28 days post-encapsulation in the samples cultured in chondrogenic medium. The samples were cultured in proliferation medium for 72 h post-encapsulation before being transferred into chondrogenic differentiation medium for an additional 25 days of culture. Representative images of the ECM markers (red) with DAPI counterstaining (blue) to localize the human ASCs, showing enhanced ECM staining for collagen I, collagen II and fibronectin within the ALG+DIM and ALG+DOM beads relative to the ALG alone beads. Abbreviations: Coll I=collagen type I, Coll II=collagen type II, FN=fibronectin. Scale bar= 200 μm.

## 4. Discussion

There remains a critical need for new treatment strategies that can promote functional meniscus regeneration. Currently, standard interventions for meniscal tears focus on short-term pain management or invasive surgical procedures if the damage is severe, which can have variable success and are associated with complications including the development of OA [1]. Tissue-engineering strategies applying scaffolds incorporating decellularized meniscus ECM represent a promising therapeutic approach [9]. An increasing body of research supports that these bioactive scaffolds can promote tissue repair by providing biological cues that can direct cell proliferation, migration, and/or differentiation, which may help to augment meniscal regeneration [13,28].

The majority of meniscal decellularization protocols reported in the literature have relied on ionic detergents and acids [14,16,17,29], which can degrade collagens and GAGs, affecting their structure and bioactivity [30]. In this study, we developed a new meniscal decellularization protocol using the milder non-ionic detergent Triton X-100 in combination with freeze-thaw and osmotic cell lysis, as well as enzymatic digestion with DNase and RNase. Triton-X100 has been used successfully for decellularization of many other tissues, including the anterior cruciate ligament [31]. Qualitative analysis using DAPI staining combined with quantification of dsDNA content validated that our new protocol was effective at extracting cells from both the inner and outer regions of the porcine meniscus. The broad combination of treatments, along with the extended treatment time, may have contributed to the efficacy of the protocol [18]. Freeze-thawing forms ice crystals that mechanically disrupt cell membranes, ultimately causing lysis [30,32], and this process can be augmented by hypotonic solution treatment, which causes an influx of water into cells [18]. These initial steps may have rendered the nuclear content more accessible to the subsequent washing and digestion steps. In addition, the decellularization protocol incorporated EDTA to sequester ions and impede cell binding to the ECM, which may have also helped to promote cell extraction [18].

Overall, immunofluorescence staining supported that key meniscus ECM constituents were retained following decellularization. Picrosirius red staining and SEM imaging confirmed that the decellularized samples showed similar collagen structure to the native tissues in both regions. Another study using a combination of 10 days of treatment with 2% Triton-X100, followed by 6 days of incubation in 2% SDS to decellularize whole porcine menisci, similarly reported minimal changes in collagen content post decellularization [33]. Although processing with Triton X-100 is generally considered to be a gentler approach compared to treatment with anionic detergents, multiple studies have reported GAG loss from tissues following Triton-X100 treatment [34–36]. In the current study, toluidine blue staining confirmed the presence of GAGs in the decellularized samples from both regions, with the DMMB assay indicating that there was a significant reduction in GAG content in the inner region.

While the pepsin digestion used to solubilize the ECM would have substantially altered the ECM structure and composition, the findings of the *in vitro* studies support that the pepsin-digested decellularized meniscus had bioactive effects on the ASCs encapsulated within the alginate beads [37]. Notably, peptides derived from the proteolytic digestion of ECM proteins, termed matrikines, are well-known to have biological activity and can modulate cell functions including growth, proliferation and differentiation [13]. High cell viability was maintained in all groups after 28 days of culture in either proliferation or chondrogenic differentiation media. Another study similarly reported high cell viability in human bone marrow-derived MSCs encapsulated and cultured in polyethylene glycol (PEG) hydrogels incorporating decellularized bovine meniscus over 7 days [38]. In addition, our analyses revealed that there was a significantly higher density of viable ASCs in the ALG+DIM and ALG+DOM beads as compared to the alginate-only controls after 28 days of culture in both media conditions. Furthermore, for the samples cultured in proliferation medium, the density of ASCs within the ALG+DIM beads was significantly higher at 28 days as compared to 24 hours.

Taken together, these results suggest that the incorporation of the digested ECM promoted cell survival and/or proliferation, or provided adhesive cues that supported cell retention within the beads. Supporting the capacity of meniscal ECM to promote cell proliferation, previous research demonstrated expansion of rat bone marrow-derived MSCs on scaffolds derived from rat meniscus, with scaffold repopulation reported within 1 week [39]. Applying a different ECM source, we have previously shown that the incorporation of decellularized adipose tissue particles can similarly promote human ASC proliferation within methacrylated chondroitin sulphate hydrogels [40].

In terms of the chondro-inductive properties of the ECM, the RT-qPCR results revealed that the expression of *SOX9* was enhanced in the 7-day samples cultured in proliferation media, suggesting that culturing within the alginate beads had an inductive effect on expression of this key transcription factor [41]. However, the generally low levels of expression of the fibrochondrogenic markers detected in the samples maintained in proliferation medium indicate that the 3D culture platforms were insufficient on their own to induce a robust chondrogenic response in the human ASCs. This response may be related to our use of passage 4 ASCs, as we have previously found that the chondrogenic differentiation of expanded human ASCs can be limited [42]. Interestingly, a previous study reported positive effects of 3D culture in proliferation medium on subsequent *in vivo* cartilage regeneration using collagen-based scaffolds seeded with auricular chondrocytes [43]. These findings support the future investigation of the chondro-inductive effects of scaffolds incorporating DIM and DOM using alternative cell populations that may have enhanced chondrogenic potential compared to ASCs, such as primary chondrocytes [43] or bone marrow-derived MSCs [43–47].

In contrast to the samples maintained in proliferation medium, culturing in differentiation medium consistently upregulated fibrochondrogenic gene expression in the human ASCs encapsulated in the alginate beads, with positive effects observed with ECM incorporation. Moreover, the immunofluorescence staining results show markedly enhanced ECM accumulation in the composite beads relative to the alginate alone after 28 days of culture in chondrogenic differentiation medium. The differing ECM composition within the two meniscal regions may have contributed to the varying effects observed between the and DOM groups, with our results suggesting that the DOM had more potent effects at stimulating the fibrochondrogenic differentiation of the ASCs. One potential mediator that was qualitatively more highly expressed in the DOM was fibronectin, which is known to play important roles in the initiation of chondrogenesis, as well as in the organization of the developing cartilage matrix [48,49]. As such, an enhanced presence of fibronectin-derived peptides within the pepsin-digested DOM may have stimulated the production of fibrocartilaginous ECM by the encapsulated ASCs. In the future, it would be interesting to apply unbiased quantitative mass spectrometry techniques to broadly compare the compositional differences between the DIM versus DOM groups, to identify other potential mediators.

Our study is not the first to report region-specific effects of decellularized meniscus. In contrast to our findings, a previous study developing bioinks reported that the incorporation of pepsin-digested decellularized porcine outer meniscus in gelatin methacryloyl/hyaluronic acid methacryloyl hydrogels enhanced collagen type I gene expression in cultured rabbit synovium-derived MSCs, while the incorporation of pepsin-digested decellularized porcine inner meniscus enhanced gene expression of markers including aggrecan and collagen type II [22]. Similarly, another study reported that the incorporation of ECM from decellularized bovine inner meniscus enhanced the fibrocartilaginous differentiation of human bone marrow-derived MSCs within polyethylene glycol diacrylate hydrogels based on collagen type II and aggrecan expression, while ECM from the outer meniscus promoted a more fibroblastic phenotype associated with collagen type I expression [38]. Many factors could have influenced these differential responses, including the decellularization methods employed, the specific cell/ECM sources studied, differences in the composition and biomechanical properties of the base hydrogels used, the ECM concentration incorporated, and differences in the methods of analysis and timepoints.

Regardless, these studies support the further investigation of the region-specific effects of decellularized meniscus to develop optimized bioinks for tissue-engineering applications that can augment cell-mediated meniscus regeneration.

## 5. Conclusions

In conclusion, a novel protocol was developed to decellularize the inner and outer regions of porcine meniscus that effectively extracted cellular content while preserving key ECM components including collagens, GAGs, and fibronectin. Taken together, the *in vitro* studies provide evidence that the incorporation of the pepsin-digested ECM within the alginate had bioactive effects on promoting cell survival, retention, and/or proliferation. Furthermore, the incorporation of the meniscus ECM enhanced fibrochondrogenic marker expression in the human ASCs encapsulated in alginate and cultured in chondrogenic differentiation medium. Gene expression and immunofluorescence staining results suggested the fibrochondrogenic differentiation response may be enhanced specifically in the ALG+DOM group, confirming that the pepsin-digested decellularized meniscus showed region-dependent effects within the hydrogels. Building from this work, future studies could investigate the effects of varying ECM concentrations within the beads, comparing the bioactivity of ECM digested with pepsin for various amounts of time [23], and assessing their potential as a bioink for 3D printing applications. Overall, this body of work contributes new insight into a growing body of evidence that supports that region-specific meniscus ECM can be harnessed to direct cell phenotype and function within tissue-engineered scaffolds.

## Acknowledgments

Funding for this study was provided by NSERC (Engage 537360-2018 and RGPIN-2017-04103), with scholarship support for S.D. and A.K from the CONNECT! NSERC CREATE Training Program and for S.D., X.Z. and A.K. from Transdisciplinary Bone & Joint Training Awards from the Bone and Joint Institute at Western University. We gratefully acknowledge Aspect Biosystems Ltd. for their generous provision of the alginate and crosslinker used in these studies. We thank our clinical collaborators Dr. A. Grant and Dr. B. Evans for their assistance with providing the adipose tissue samples used for cell isolation in these studies. Finally, we thank Dr. Todd Simpson for his assistance with the SEM imaging, which was performed at the Western Nanofabrication Facility.

## Ethics Approval Statement

Human research ethics board approval was obtained from Western University for the studies involving human adipose tissue (HSREB #105426), and all tissues were obtained with informed consent.

## Data Availability Statement

The raw data supporting the conclusions of this manuscript will be made available by the authors, without undue reservation, to any qualified researcher.

