## Supplementary Figures for "Investigation of region-specific effects of pepsin-digested decellularized meniscus on human adipose-derived stromal cells within alginate hydrogels"

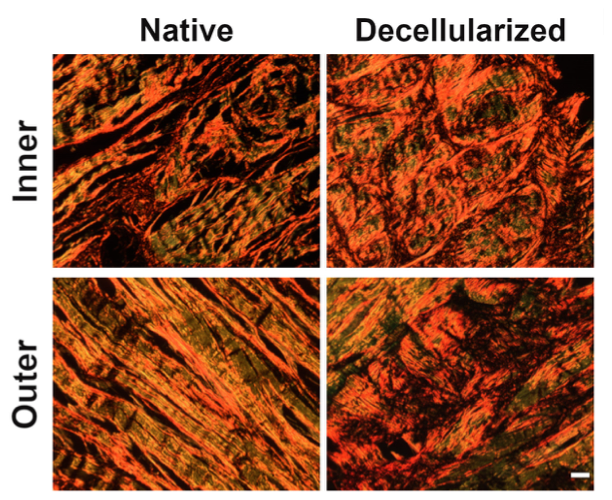

**Supplemental Figure 1. Pircosirius red staining patterns remained similar following decellularization.** Representative staining with visualization using polarized light microscopy showing that the decellularized tissues contained a dense network of collagen fibers, with similar staining patterns to the native tissue samples in both regions. Scale bar = 200 µm.

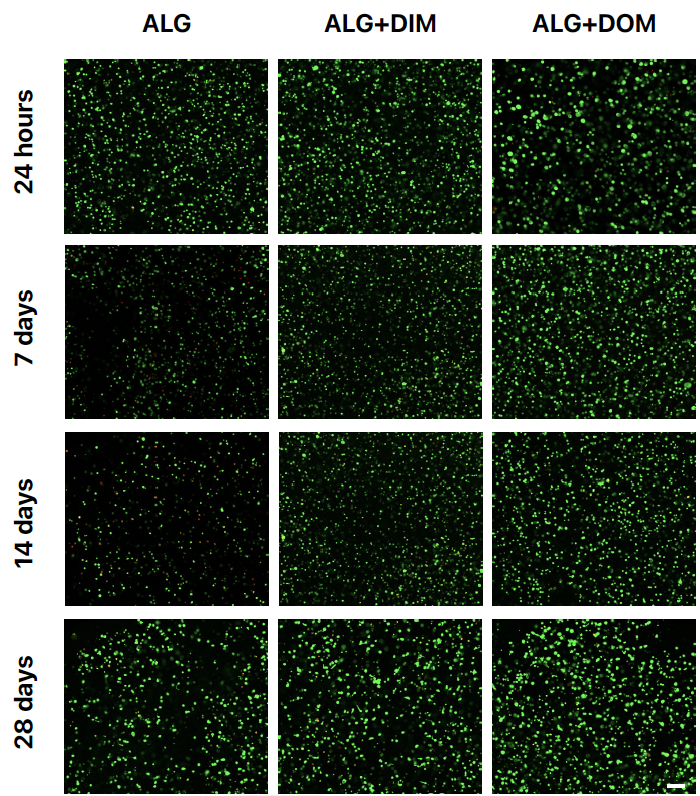

**Supplemental Figure 2. LIVE/DEAD confirmed that the human ASCs remained highly viable over 28 days following alginate bead encapsulation culture in proliferation medium.** Representative images showing calcein+ live (green) and EthD-1+ dead (red) ASCs in the ALG, ALG+DIM, and ALG+DOM beads in the samples cultured in proliferation medium. Scale bar = 100 μm.

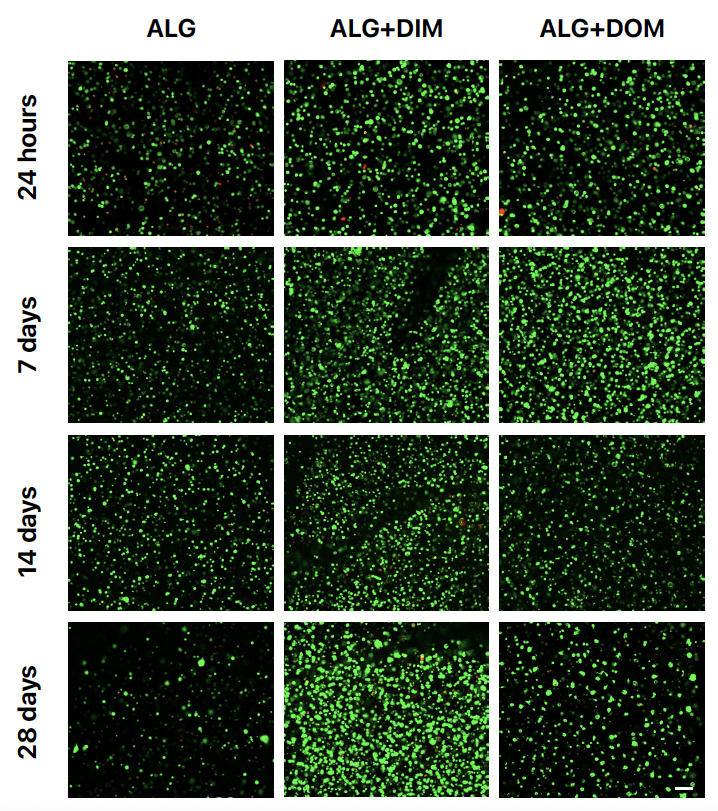

**Supplemental Figure 3. LIVE/DEAD confirmed that the human ASCs remained highly viable over 28 days following alginate bead encapsulation and culture in chondrogenic differentiation medium.** The samples were encapsulated and cultured for 3 days in proliferation medium, prior to be transferred into chondrogenic differentiation medium. Representative images showing calcein+ live (green) and EthD-1+ dead (red) ASCs in the ALG, ALG+DIM, and ALG+DOM beads at time points post-encapsulation. Scale bar = 100 μm.

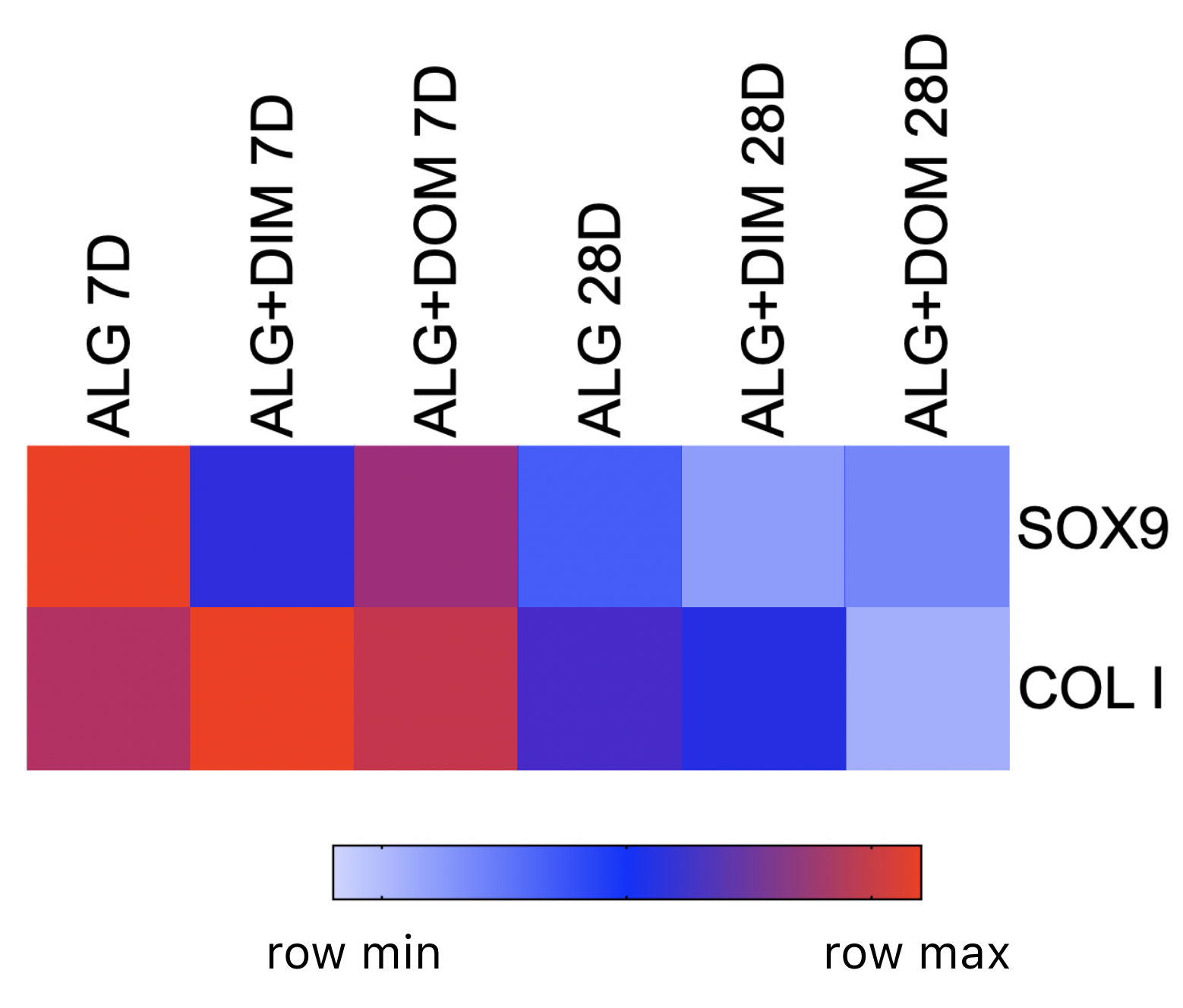

**Supplemental Figure 4. Consistent gene expression patterns in human ASCs encapsulated within the alginate-based beads and cultured in proliferation medium over 28 days.** *COMP*, *ACAN*, *Coll II*, and *Coll X* were excluded due to undetected or inconsistent low levels of expression between the two cell donors. Data was analyzed using the ∆∆Ct method with normalization to the geometric mean of the stable housekeeping genes *RPL13A* and *GUSB*, and using the 7-day alginate beads cultured in chondrogenic differentiation medium as the calibrator. The heat map represents the average relative gene expression for the two cell donors, with similar patterns observed between the donors. (n=3 bead samples processed separately at each timepoint/trial, N=2 trials with different ASC donors).

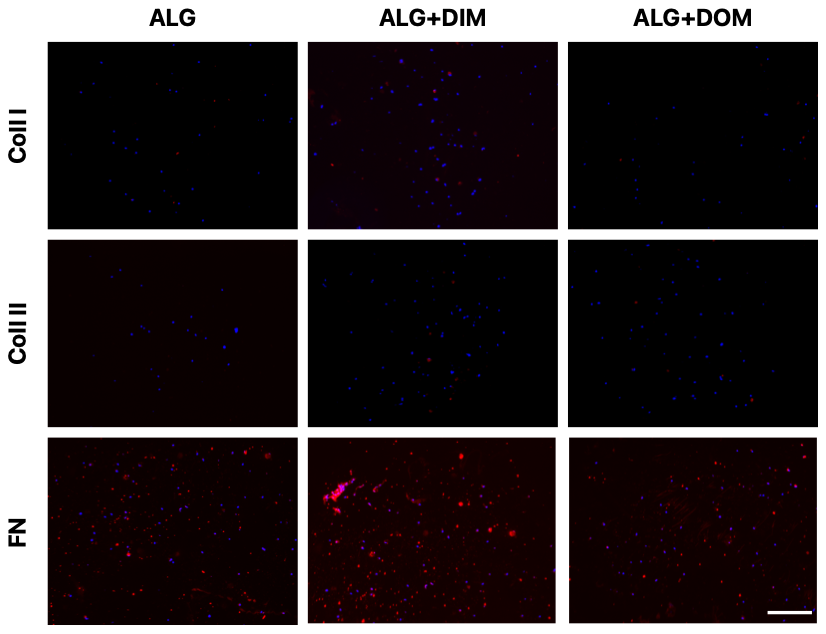

**Supplemental Figure 5. Immunohistochemical staining showed limited ECM marker expression within the alginate-based beads following 28 days of culture in proliferation medium.** Representative images of the ECM markers (red) with DAPI counterstaining (blue) to localize the human ASCs. Fibronectin expression was qualitatively enhanced in the ALG+DIM group. (n=3 cross-sections containing multiple beads/trial, N=2 trials with different ASC donors). Abbreviations: Coll I=collagen type I, Coll II=collagen type II, FN=fibronectin. Scale bar= 200 μm.
